# Leveraging the p53 signaling pathway as a radio-sensitization strategy in endometrial cancer

**DOI:** 10.1101/2022.09.06.505009

**Authors:** Roberto Vargas, Aaron Petty, Brian Yard, Arda Durmaz, Kristi Lin-Rahardja, Ofer Reizes, Robert Debernardo, Jacob Scott

## Abstract

Endometrial cancer (EC) is the most common type of gynecologic malignancy in the United States, with over 66, 200 new cases expected in 2023. The number of mortalities per year now approximate that of ovarian cancer. Despite our ability to identify different biologic clusters of EC, we have yet to understand the functional impact of key genomic alterations associated with discrepant prognoses and exploit this knowledge for therapeutic benefit. Given this, we set out to determine how alterations in p53 signaling, as conferred my *TP53* mutations, impact radiotherapy response in EC. We also explored if manipulation of this signaling pathway could be utilized as a radio-sensitization strategy in EC. Our work demonstrates that p53 signaling plays a significant role in radiotherapy response for EC and that leveraging this genomic data may allow us to exploit this pathway as a viable radiotherapeutic target in a significant number of EC cases.

## Introduction

Endometrial cancer (EC) is the most common gynecologic malignancy in the United States, with over 66, 200 new cases expected in 2023.^1^ Current trends outpace prior projections, with a 55% increase in incidence predicted from 2010 to 2030.^2^ Endometrial cancer mortality rates now approach that of ovarian cancer, highlighting the need for increased investigative focus. There have been a significant advance in determining the genomic underpinnings of EC ^3–7^ and development of prognostic capabilities based on these findings.^8,9^ Despite our ability to identify different biologic clusters of EC, we have yet to fully understand the functional and therapeutic impact of the key genomic alterations associated with prognosis groups, such as *TP53* mutated EC, and exploit this knowledge to provide innovative treatment strategies.

Historically radiotherapy has played a key role in the prevention and treatment of disease recurrence for both early-stage and advanced EC.^7,10–12^ Recent clinical trials have challenged the indications for radiotherapy in EC, particularly in those patients with high-risk *TP53*-mutated cancers.^13,14^ The PORTEC-3 study demonstrated that the addition of chemotherapy improved outcomes in those tumors of serous histology. A subsequent molecular study demonstrated that this benefit extended to all tumors with aberrant p53, as identified via immunohistochemistry (IHC).^15^ The results of the GOG-258 provided additional clinical data challenging the value of radiotherapy entirely, with chemo-radiation providing no survival benefit over chemotherapy alone. While these two landmark studies provide evidence that introduces uncertainty behind long-standing paradigms, they nonetheless still highlight the important role radiotherapy plays in reducing pelvic and vaginal recurrences. They also introduce the concept of tailoring EC treatment based on histologic or genomic findings, recognizing that *TP53* mutated tumors benefit from treatment escalation.^14^ Despite these robust clinical observations, the functional impact on TP53 on radiotherapy response has not been explored in EC.

Conflicting information exists regarding the role of p53 signaling and its tissue-specific impact on radiotherapy response^16,17^.^18^ Differential response to radiotherapy is observed in human tissues despite similar expression levels of wild-type p53, indicating that other tissue-specific factors may impact the role this pathway plays on radiotherapy response^19,20^. The lack of growth-arrest associated with aberrant p53/p21 signaling has been tied to both radiation resistance^21^ and sensitivity.^22^ Studies interrogating the role of mutant p53 on radiation response have also led to inconsistent results, with some demonstrating no effect on radiation response^23,24^, while others note a correlation with radiation resistance^25^ ^26^ The lack of robust experimental data specific to EC, compounded by the dualistic role presented in the literature dampen our ability to attribute poor clinical outcomes in *TP53* mutated EC to radio-resistance.

Current clinical EC studies recognize aberrant p53 immunohistochemical (IHC) staining as a clinically relevant pathologic marker. Both absent IHC staining and “strong” staining are considered abnormal, with the former correlating with loss the protein entirely and the latter representing the accumulation of mutated p53 protein. Despite the clinical utility of this stain in broadly identifying EC with abnormal *TP53,* it does not allow for consideration of allelic frequencies and any remaining wild-type/functional copies of the gene.^27,28^ Allelic frequency of *TP53* mutations has previously been demonstrated to differentially impact clinical outcomes in myelodysplastic disorders^29–33^ and contribute to the development of somatic copy number alterations^34^. This functional and clinical observations highlight the potential to employ more precise, clinically-meaningful genomic data to stratify EC outcomes. Understanding the functional impact of allelic frequency, serving as a proxy for residual wild-type p53 function, may allow for personalization of radiotherapy based on next-generation sequencing data.

The increased prevalence in this understudied malignancy, coupled with the limited functional data to account for the worse outcomes in p53-aberrant EC, prompted our investigation into the association between p53 signaling and radiotherapy response in endometrial cancer. To accomplish these goals, we integrated a validated high-throughput radiation phenotyping platform, intron-targeting CRISPR/Cas9 techniques, and dose-response assessment using cell lines. Additionally, we evaluated the role of pharmacologic manipulation of this signaling pathway using MDM-2 inhibitors, as a novel radio-sensitization strategy for EC. Lastly, we queried the TCGA database to assess for the potential benefit of allelic frequency stratification in *TP53*-mutated EC patients. We hope that our findings provide the biologic rationale to support the findings of published clinical trials and the continued application of radiotherapy for clinical benefit, with increased focus on precision radiotherapy leveraging functional genomics.

## Results

### *TP53* status impacts radiotherapy response in EC cell lines

The examine the association of *TP53* mutations and their allelic frequencies on radiation resistance we correlated *TP53* status with gamma-radiation response in EC cell lines, using area under the curve (AUC) measure from a pan-cancer profiling effort. ^35^ Endometrial cancer cell lines with altered *TP53* status were more likely to have higher radiation resistance/area under the curve (AUC) (LR 11.12, p = 0.004). Given the potential impact of allelic frequency on therapeutic response, we then annotated the analysis based on allelic frequencies reported in Cancer Cell Line Encyclopedia.^36^ When stratified by allelic frequency, those with a VAF >0.5 (high VAF) were also more likely to be associated with radiation resistance/higher AUC (LR 9.42, p=0.009), as seen in **Figure 1A**. Using an intron-targeting CRISPR/Cas9 system, which spares lentiviral constructs lacking intronic segments, we then conducted a complementation experiment with a subset of EC cell lines. A statistically significant increase in the AUC (representative of radiation resistance) was observed when JHUEM-1 and JHUEM-2 cell lines (both wild-type *TP53)* underwent CRISPR/Cas9 knock-out (KO) of *TP53* (**Figure 1B**). As noted on Western blotting, signaling through the p53/p21 pathway was disrupted with KO (**Figure 1C**). No discernible change in the AUC for radiation response was observed in CRISPR/Cas9 *TP53* knockouts in the Hec1A, Hec1B, and MFE-319 cell lines, all of which contained high-VAF GOF/DN *TP53* alterations (**Supplementary Figure 1**).

**Figure 1.**
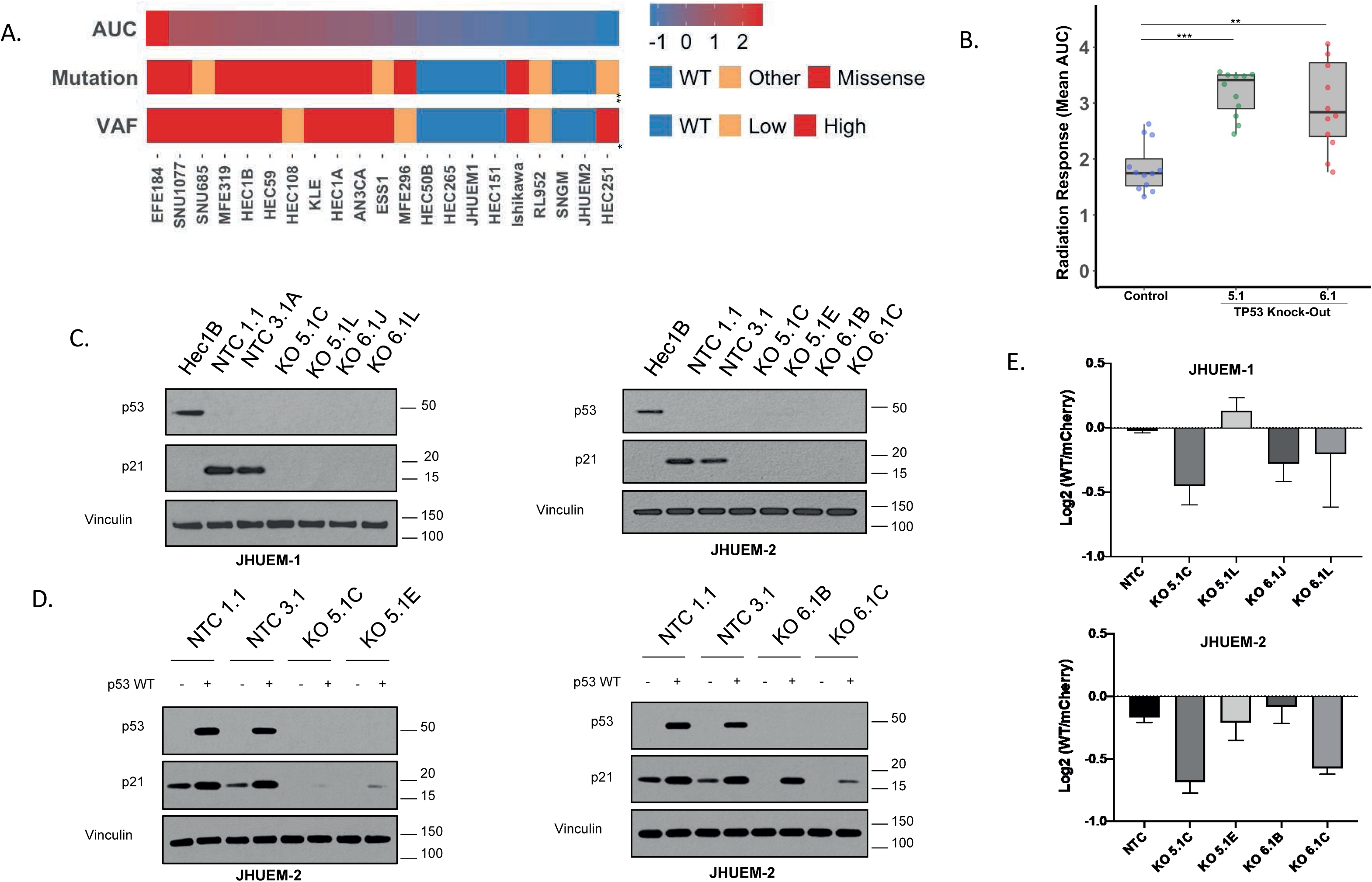
*TP53* Status and AF are Associated with Radiation Resistance in EC Cell Lines. A) Heat-map depicting 21 EC cell lines organized by decreasing radiation resistance (left to right) as quantified by area under the curve (AUC). The cell lines’ *TP53* mutational status and VAF are depicted in the bars below, denoting the significant associations between AUC and *TP53* mutation status (p = 0.004) or VAF (p = 0.01), respectively. A linear regression model was fitted to assess for the correlation between mutation and AUC (Likelihood Ratio = 11.12 and P= 0.0038) as well as VAF and AUC (Likelihood Ratio = 9.42 and P= 0.009). B) Box-whisker plot depicting the increase in radiation AUC noted in JHUEM1 and JHUEM2, using two different CRISPR/Cas9 guide-RNA (gRNA) sequences for *TP53* compared to the non-targeting control (NTC). Kruskal-Wallis test with Dunn’s Multiple Comparisons Test was performed with p< 0.001 for NTC vs KO 5.1, p = 0.0016 for NTC vs KO 6.1. C) Western-blot analysis of two monoclonal isolates from each gRNA and NTC reveals loss of p21 signaling with knock-out of *TP53*. Hec1B is our positive control for p53. D-E) Complementation with wild-type *TP53* re-establishes p21 signaling and radiation sensitivity in knock-out clones of JHUEM-2. For sub-figure E and JHUEM2, one-way ANOVA test with Dunnett’s Multiple Comparisons test was performed with p = 0.0051 for KO 5.1C vs NTC, p = 0.0199 for KO 6.1C vs NTC.

Complementation with a low-expression (PGK) vector containing wild-type *TP53* re-established p53/p21 signaling (**Figure 1D**) and radiation sensitivity (Figure 1E) in JHUEM-1/JHUEM-2 that had parental wild-type alleles previously knocked-out. Complementation of our copy-number high/*TP53-*KO constructs (Hec1A, Hec1B, and MFE319) with a wild-type *TP53* vector did not reintroduce significant radiation sensitivity. It is plausible that the evolutionary trajectories of these *TP53-*mutant cell lines have allowed for the amplification of other established genomic drivers of resistance, such as somatic copy-number alterations, aside from the direct impacts of p53/p21 signaling.

### Gain-of-function/dominant-negative *TP53* mutations negatively impact radiotherapy response

Employing the same *TP53-KO* constructs we subsequently explored the impact of five hotspot *TP53* alterations (R248Q, R248W, R273C, R273H, Y220C) in EC.^27,28,37^ First, the effect of each mutation on p53/p21 signaling and radiation phenotype was assessed. We noted that the introduction of these hotspots into a *TP53-*KO (p53 null) cell line did not re-establish p53/p21 signaling (**Figure 2A**, **Supplementary Figure 2**) or change radiation resistance. When the mutations were co-expressed with wild-type *TP53* present in our non-targeting controls for both JHUEM-1 and JHUEM-2, a dominant-negative effect was confirmed. The 5 hotspot variants led to an increase in radiation resistance, despite the presence of two parental wild-type alleles (**Figure 2B, Supplementary Data**). Concurrently, an abrogation of the p53/p21 signaling was observed **(Figure 2C**), supporting the notion that p53 signaling through the p21 pathway is negatively affected by these variants and contributes to resistance.

**Figure 2.**
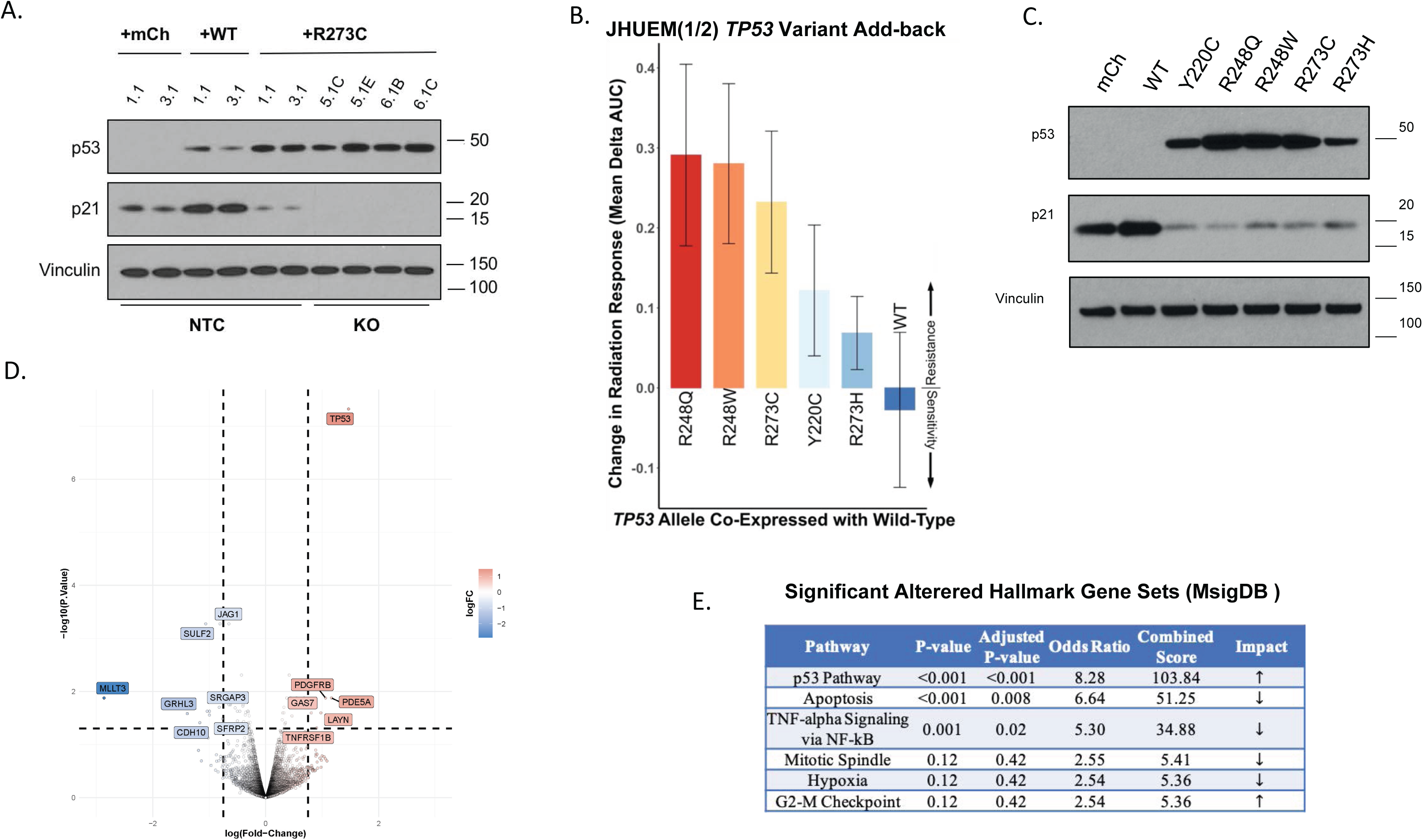
Gain-of-Function/Dominant-Negative Alleles of *TP53* Impact p21 Signaling, Radiation Response, and Apoptotic Pathways. A) Western-blot analysis of JHUEM-2 non-targeting control [NTC] isolates (1.1 and 3.1) with the addition of mcherry (complementation transfection control), wild-type *TP53* (over-expression), and the addition of the R273C allele (co-expression with wild-type). A decrease in p21 signaling after introduction of the R273C allele is noted in the NTC isolated, despite parental/wild-type *TP53* alleles. In the knockout isolates (5.1C/E and 6.1B/C) there is no p21 signaling observed with the mutant allele alone. B) Waterfall plot depicting the overall effect on AUC after introduction of a variant allele (or wild-type) across 4 different mono-clonal NTC isolates. As noted, introduction of variant alleles led to an increase in radiation resistance from baseline despite the presence of a wild-type allele. C) Western-blot analysis demonstrating decrease in p21 signaling with each analyzed GOF/DN *TP53* variant. D) Volcano plot analysis of an the NTC isolates with R273C co-expression, compared to the NTC alone (wild-type) 24hr after 2Gy irradiation. An increase in the expression of *TP53* in our wt/R273C isolate confirms increased expression of p53, with a decrease in the expression of genes associated with apoptosis and TNF-alpha signaling (E, Supplementary Table 2).

To explore for potential off-target effects of dominant-negative variants, we treated our JHUEM2 non-targeting controls (NTC) with parental wild-type alleles and the JHUEM2 (NTC + R273C) constructs with 2Gy of gamma radiation. RNA was extracted 24 hours later and differential gene expression analysis was performed. Hallmark pathways that were significantly altered in the *R273C* co-expressing cell lines (1.1RC and 3.1RC) included apoptosis and tumor necrosis-alpha gene sets, mainly driven by down-regulation of *CDKN1A, BTG2, CCND1, GADD45B, NINJ1, IER2, and FAS* (**Figure 2D-E, Supplementary Figure 3, Supplementary Table 1**).

### MDM-2 inhibition sensitizes EC cell lines by increasing wild-type signaling

MDM-2 inhibitors are a class of small molecules that inhibit the E3-ubiquitin ligase responsible for the degradation of p53. MDM-2 inhibitors have been tested in other solid malignancies, but their role in the treatment of EC has not been assessed. Given the significance of the p53/p21 signaling pathway in EC cell lines, we tested the hypothesis that MDM-2 inhibitors would provide a cytotoxic and/or radio-sensitizing strategy in EC. Nutlin-3 is a first-generation MDM-2 inhibitor, while AMG-232 is a potent MDM-2 inhibitor undergoing clinical trial testing in soft-tissue sarcomas.^38^ Using three parental cell lines to replicate a wild-type (JHUEM2), low-VAF (Hec 108), and high-VAF (Hec1B) endometrial malignancy, we performed cell viability experiments using both Nutlin-3 and AMG-232. A differential effect on cell viability was observed depending on the presence of wild-type *TP53,* co-expression of wild-type/variant, or expression of only the GOF/DN allele. Only the cell lines with a wild-type allele present (JHUEM2 and Hec108) have decreased cell viability (**Figure 3A**) and downstream p21 activation (**Figure 3B**) in response to treatment with Nutlin-3. Treatment of three parental EC cell lines with AMG-232 confirmed the more potent effect on cell viability (**Figure 3C**) and induction of p21 (**Figure 3D**) than with the first-generation MDM-2 inhibitor. Hec1B, with high-VAF *TP53,* is resistant to treatment with both MDM-2 inhibitors, consistent with prior studies.^39^ To assess clinical relevance, we queried the TCGA cohort for MDM2 expression in EC. We noted a negative correlation between MDM2 expression and *TP53* status; patients with wild-type or low-VAF *TP53* mutant EC retained some expression of MDM2 (**Figure 3E**).

**Figure 3.**
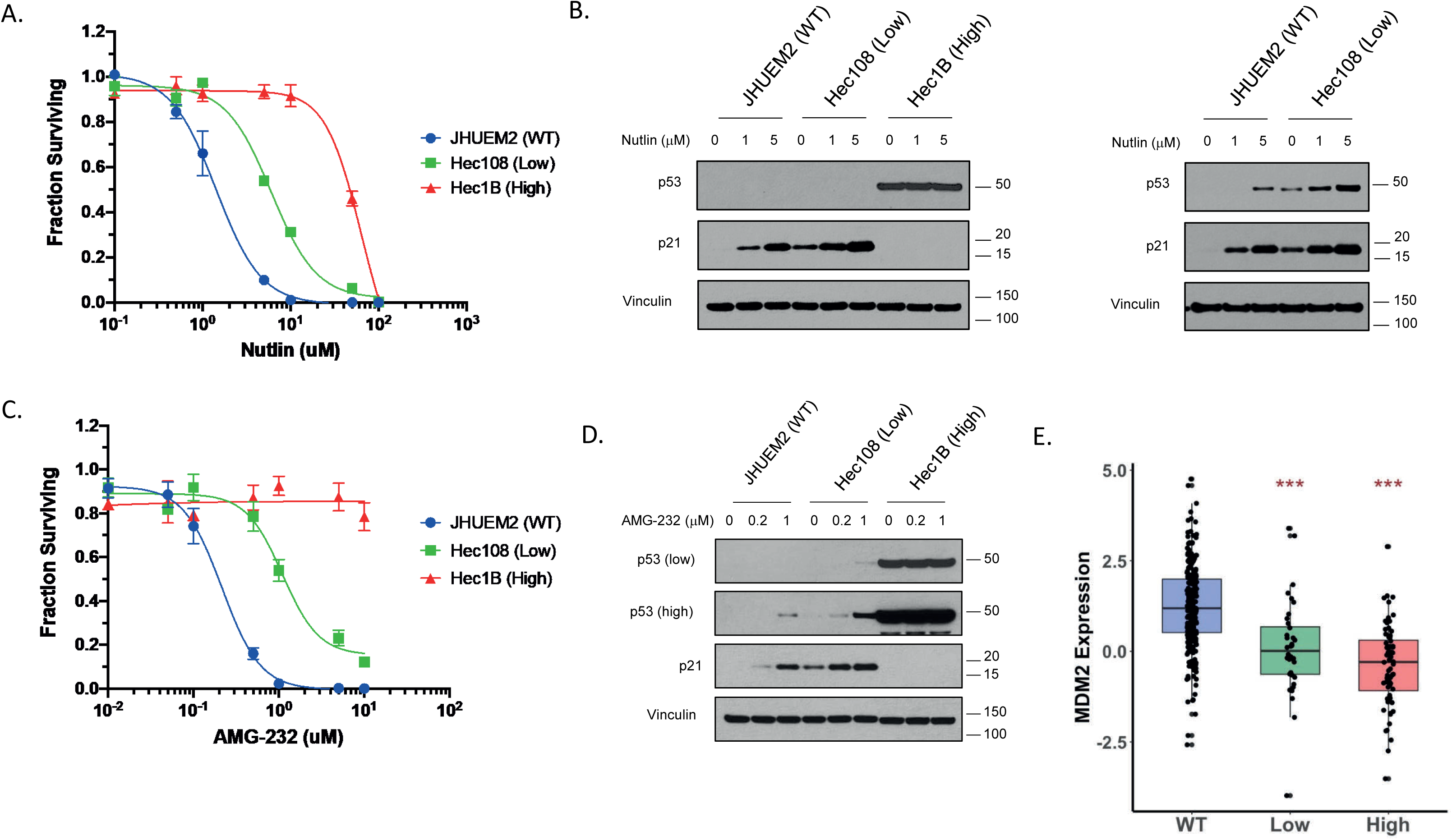
MDM2 Inhibition with Nutlin-3 and AMG-232 as a Cytotoxic Strategy. A, C) Drug response curve of EC cell lines representing three different VAF populations – JHUEM2 (wild-type), Hec108 (low-VAF), and Hec1B (high-VAF). Nutlin-3 (A) and AMG-232 (C) both demonstrate significant effects on cell viability in JHUEM2 and Hec 108. Hec1B was resistant to treatment with either drug. B, D) Western-blot analysis of all three cell lines demonstrate an increase in p53/p21 with increasing concentrations of either drug. E) Box-whisker plot of MDM2 expression across TCGA patients demonstrate that MDM2 expression decreases with increasing VAF, but there remains a significant portion of low-VAF patients with mRNA expression that could be targeted via Nutlin-3/AMG-232. For subfigure E, one-way ANOVA with Tukey’s Multiple Comparisons Test was conducted to analyze relative expression compared to the WT group. p < 0.001 for Low vs WT; p < 0.001 for High vs WT.

When the same cell lines are treated with Nutlin-3 and exposed to a range of doses of gamma radiation, a synergistic effect is observed in JHUEM2 and an additive effect in Hec 108 (**Figure 4A, 4C**). AMG-232 showed a synergistic effect in both the JHUEM2 and Hec108 *TP53*-mutant cell lines (**Figure 4B, 4C, Supplementary Figure 4**). These findings deepen our understanding of the relationship between MDM-2 inhibition and *TP53*, allowing for potential therapeutic development and novel applications in EC.

**Figure 4.**
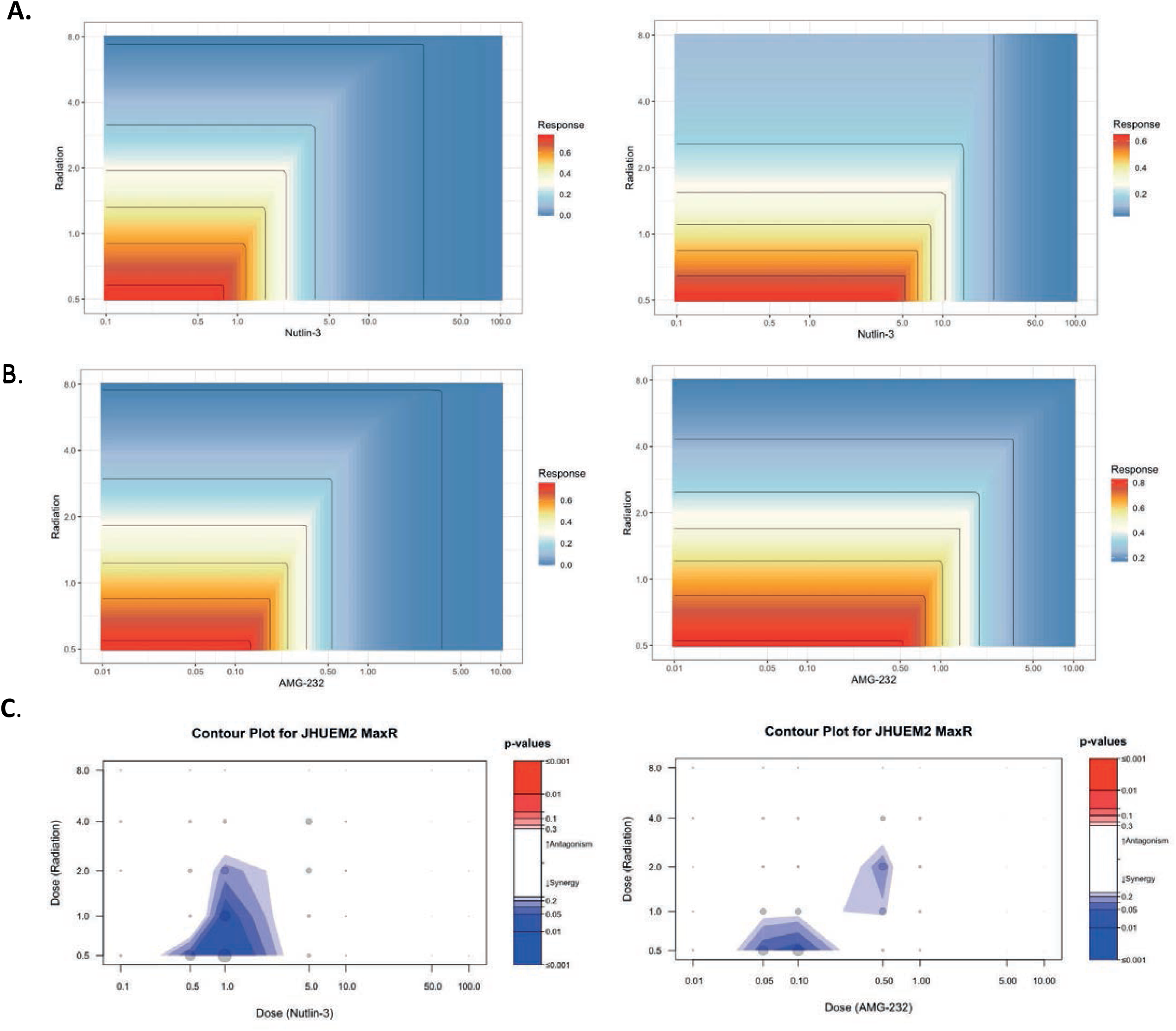
MDM2 Inhibition Is Synergistic with Radiotherapy Dependent on *TP53* Status. A-B) Isobolograms for Nutlin-3 (A) and AMG-232 (B) in two EC cell lines - JHUEM2 (left) and Hec108 (right) - treated with gamma irradiation. As depicted, increasing drug dosage is the x-axis and increasing radiation dose is the y-axis. A robust effect on Nutlin-3 and AMG-232 is achieved at low drug concentrations and low RT doses for both the wild-type (JHUEM2) and low VAF (Hec108) cell line. C) Contour plots quantitatively demonstrate the synergistic effect of Nutlin-3 in JHUEM2 (left), as well as with AMG-232 in JHUEM2 (right) and Hec108(Supplementary Figure 5). The remaining interactions were classified as additive. Synergy was quantified by comparing actual values to those predicted by fitting the Highest Single Agent (HSA) model to the data.

### *TP53* variant allelic frequency is associated with worse clinical outcomes in EC

To assess for the potential clinical impact of TP53 mutations and allelic frequency on radiotherapy response for EC, we queried the TCGA dataset.^37^ Data from comprehensive genomic profiling effort was carefully annotated to include only state I-III EC patients and further annotated for those coded to have received radiotherapy. Specific dosing and route of administration of radiotherapy could not be accounted for in repository. We also excluded patients with stage IVB disease or those annotated to have received chemotherapy, to restrict our observations to radiotherapy response alone. We found that the prevalence of *TP53* mutations increased with increasing sCNA burden, but not exclusively limited to that group (**Figure 5A**). Patients who received radiotherapy were more likely to have higher-grade tumors and increased tumor mutational burden, compared to those who do not receive radiotherapy (**Supplementary Table 2**). While tumor mutational burden is not clinically considered when prescribing adjuvant radiation, tumor histologic grade is a known recurrence risk factor and is often considered when determining adjuvant therapy. On univariate analysis, histologic grade, FIGO stage, SCNA, and *TP53* status were all associated with worse progression-free survival, or PFS (**Supplementary Table 2**). On multivariable analysis only the FIGO stage was associated with a worse prognosis, supporting the notion that *TP53* mutations and copy-number high status often co-exist in EC (**Supplementary Table 2**).

**Figure 5.**
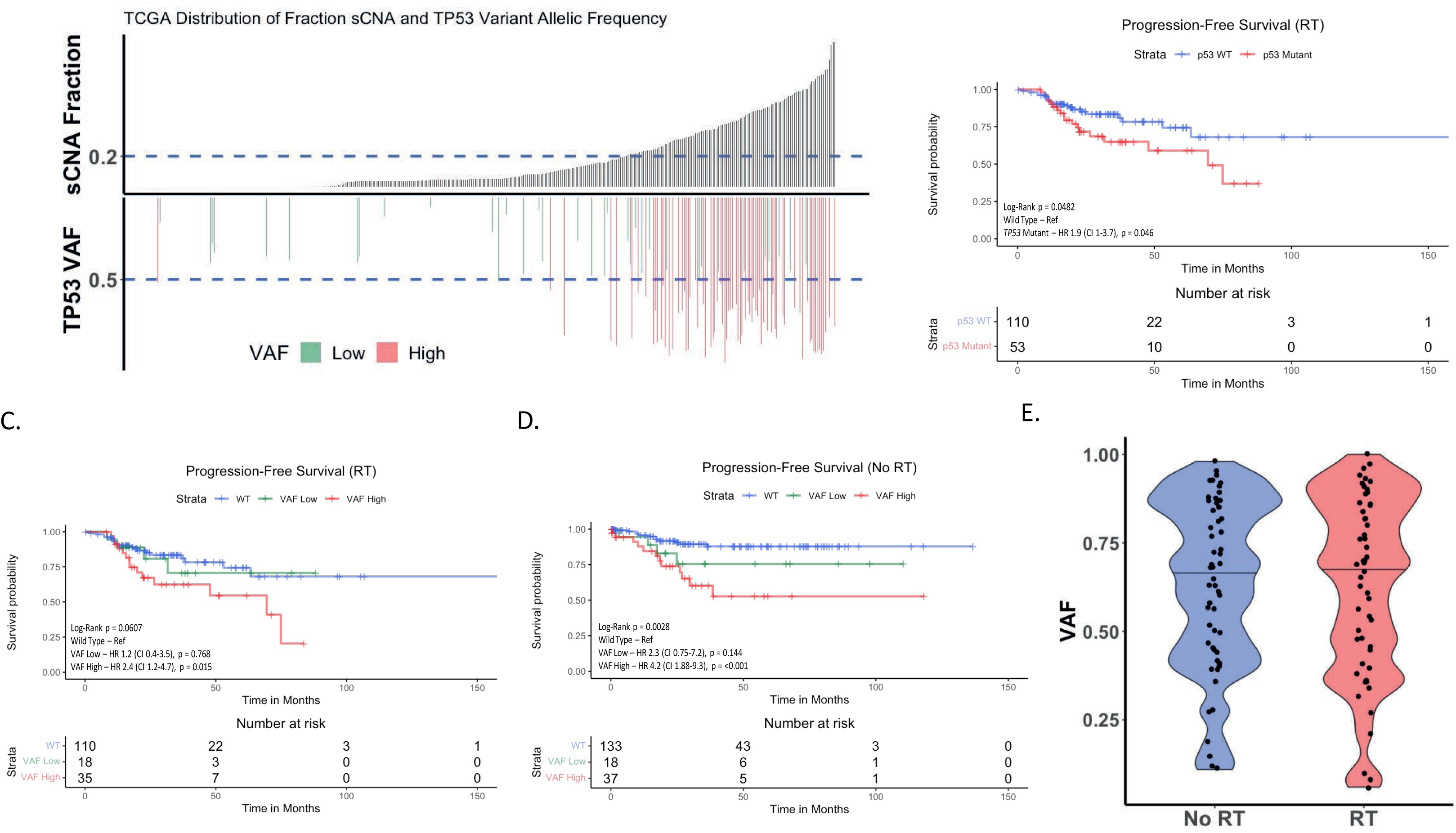
*TP53* Variant Allelic Frequency Is Clinically Relevant in Endometrial Cancer. A) This plot represents available EC patients in cbioportal with somatic copy number alteration (sCNA) burden and *TP53* data available. The dashed line at 0.2 defines the cutoff for copy-number high used in TCGA. As noted, the frequency of *TP53* mutations increases with increasing sCNA. While the majority of *TP53* mutations have a high variant allelic frequency (VAF), there remain patients with <50% allelic frequency throughout the entire population. B-D) Kaplan-Meier plots and Cox-regression for progression free survival (PFS) stratified by *TP53* status (B) and *TP53* VAF (C-D). As noted in subfigure B, patients with *TP53* mutation had worse PFS compared to wild-type patients when treated with RT. When stratified by VAF (C), patients with a high VAF had a worse PFS compared to low VAF and wild-type patients. We observed the same in non-RT patients (D), albeit the difference in risk of progression based on hazard ratio was more notable between low VAF and wild-type. E) Violin plot representing the overall distribution of *TP53* VAF in both treatment groups, highlighting that current treatment patterns do not take VAF into consideration.

Patients with *TP53* alterations had a significantly worse PFS (HR 1.9, 95% CI 1.0 - 3.7, log-rank p = 0.046) compared to patients with wild-type *TP53,* despite treatment with RT (Figure 1B). For assessment of allelic frequency, a VAF cutoff of 0.5 was employed to capture tumors with potential loss-of-heterozygosity (LOH) and thus reduced wild-type *TP53*. When stratified by *TP53* variant allelic frequency (VAF), only patients with a VAF > 0.5 (high VAF) had a significantly worse PFS after RT treatment, compared to those with *TP53* VAF <=0.5 (low VAF) (HR 2.01, 95% CI 0.66 - 6.1) and wild-type *TP53* tumors (HR 2.4, 95% CI 1.2 – 4.7, log-rank p = 0.015). In the face of radiotherapy, low VAF patients responded most like WT patients (**Figure 5C, Supplementary 5**). The differential outcome based on VAF was also observed with patients who did not receive RT, highlighting the established and known prognostic value of *TP53* status in EC. Overall the trend in the hazard ratios in low VAF patients suggests that *TP53* allelic frequency may also have predictive value in regards to radiation response (**Figure 5D and Supplementary Figure 5**), but this remains to be studied in a robust and prospective manner. While most patients in the “serous-like” cluster will have a high VAF (>0.5) *TP53* mutation, 15% of these high-risk patients retain a low VAF (Table 1) and could retain some form of wild-type function.

**Table 1.**
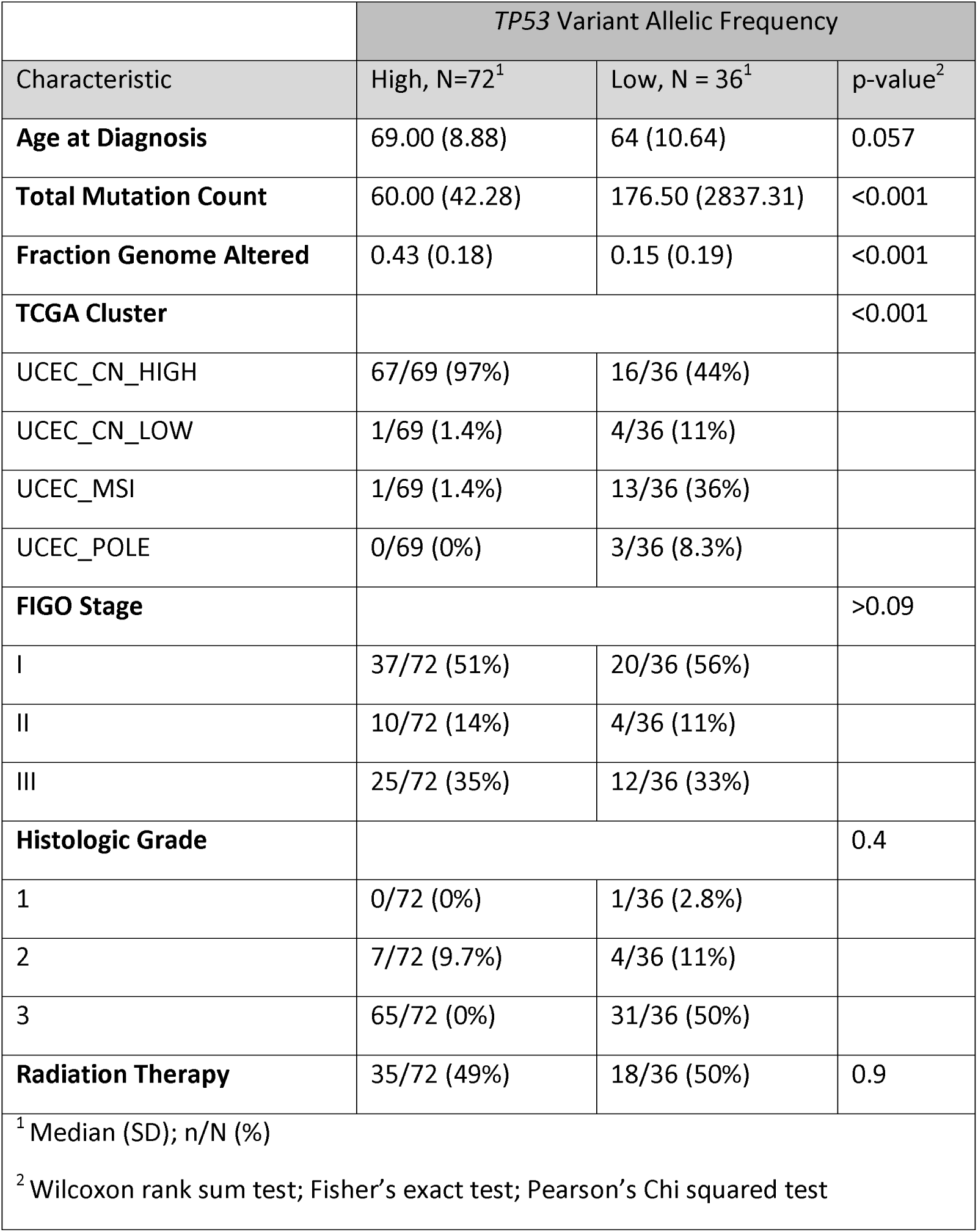
Stratification of *TP53* mutant patients based on VAF highlights other associated genomic markers. Demographics table of patients with TP53 alterations stratified by their variant allelic frequency. High is defined as >50% and low is defined as <=50%. As noted, patients with a low VAF were more likely to have higher mutational counts, lower copy-number alterations, and less likely to be classified into the “serous-like” (UCEC_CN-HIGH) TCGA cluster. When taken as an entire group, the “serous-like” cluster had 105 patients, with 16 (15%) falling into the low VAF classification. No differences in clinical stage, histologic grade, or receipt of adjuvant radiation (RT) therapy was noted.

|  | *TP53* Variant Allelic Frequency |  |  |
| --- | --- | --- | --- |
| Characteristic | High, N=72<sup>1</sup> | Low, N = 36<sup>1</sup> | p-value<sup>2</sup> |
| **Age at Diagnosis** | 69.00 (8.88) | 64 (10.64) | 0.057 |
| **Total Mutation Count** | 60.00 (42.28) | 176.50 (2837.31) | <0.001 |
| **Fraction Genome Altered** | 0.43 (0.18) | 0.15 (0.19) | <0.001 |
| **TCGA Cluster** |  |  | <0.001 |
| UCEC_CN_HIGH | 67/69 (97%) | 16/36 (44%) |  |
| UCEC_CN_LOW | 1/69 (1.4%) | 4/36 (11%) |  |
| UCEC_MSI | 1/69 (1.4%) | 13/36 (36%) |  |
| UCEC_POLE | 0/69 (0%) | 3/36 (8.3%) |  |
| **FIGO Stage** |  |  | >0.09 |
| I | 37/72 (51%) | 20/36 (56%) |  |
| II | 10/72 (14%) | 4/36 (11%) |  |
| III | 25/72 (35%) | 12/36 (33%) |  |
| **Histologic Grade** |  |  | 0.4 |
| 1 | 0/72 (0%) | 1/36 (2.8%) |  |
| 2 | 7/72 (9.7%) | 4/36 (11%) |  |
| 3 | 65/72 (0%) | 31/36 (50%) |  |
| **Radiation Therapy** | 35/72 (49%) | 18/36 (50%) | 0.9 |
| <sup>1</sup> Median (SD); n/N (%) |  |  |  |
| <sup>2</sup> Wilcoxon rank sum test; Fisher's exact test; Pearson's Chi squared test |  |  |  |

## Discussion

Our integrative analysis incorporating high-throughput radiation phenotyping, complementation experiments using CRISPR/Cas9, and pharmacologic targeting, and a clinically-annotated dataset provides a full-spectrum fundamental biologic rationale behind the disparate clinical outcomes in EC. While the prognostic value of *TP53* status in EC has previously been observed in clinical trials, the precise therapeutic impact remained poorly understood. Using cell lines, we demonstrated the oncogenic gain-of-function impact and dominant-negative effects of *TP53* variants observed in EC. We also established that not all *TP53*-mutated cell lines are resistant to MDM-2 inhibition, and the degree of resistance depends on the variation in the amount of functional wild-type p53. Lastly, we demonstrated that allelic frequency of *TP53* alterations is prognostic, and possibly predictive, of clinical outcomes in EC.

Given that the majority (80%) of EC have WT *TP53,* we first focused on wild-type/low-grade cell lines to demonstrate that the loss of this key oncogene resulted in the development of radiation resistance. Subsequently, we noted that restoration of a wild-type allele re-established downstream WAF/p21 signaling and restored a small degree of radiation sensitivity. We attribute the low expression of wild-type *TP53* by the PGK promoter in our vector to explain the lack of a complete restoration in radiation response-phenotype. Using the same complementation approach also allowed for the assessment of common hotspot mutations (Y220C, R273C/H, and R248Q/W) and their functional impact. While the mutations themselves provided a GOF phenotype by conferring radioresistance, they also exhibited a dominant-negative effect on wild-type signaling. Despite a low-expression vector, the introduction of all 5 missense mutations suppressed p21 signaling established by the two parental wild-type *TP53* alleles. Furthermore, we demonstrate that the GOF/dominant-negative alleles also conferred a radio-resistant phenotype.

Gene-expression changes observed after treatment with 2Gy of radiation support the notion that these common mutations may be targeting other pathways relevant to radiation response. Our pathway analysis shows genes associated with apoptotic pathways and tumor necrosis factor-alpha (TNFα) signaling via NF-κB are down-regulated in the presence of the R273C allele and irradiation. Pal *et al*. previously demonstrated that TNFα influences radiation-mediated DNA damage, with decreased TNFα leading to reduced double-stranded breaks in lung cancer cell lines.^40^ Additionally, the same authors found that serum TNFα expression positively correlated with clinical outcomes after radiotherapy. The degree of TNFα induction after irradiation has also been shown to predict clinical radiotherapy response in multiple solid tumors.^41^ While TNFα/NF-κB signaling has been shown to have both pro-apoptotic and anti-apoptotic properties based on cell/tumor lineage^42^, in our *TP53-*altered (R273C) EC cell line we saw a decrease in the expression of genes associated with TNFα/ NF-κB signaling, as well as other apoptosis-related genes when treated with radiation. Prior studies in prostate cancer cell lines have also noted a decrease in NF-κB is associated with anti-apoptotic effects after radiation.^43,44^ Crosstalk between p53 and TNFα/NF-κB through non-canonical pathways may be contributing to cell survival after irradiation.^45^

This ongoing need for novel treatment strategies incorporating radiotherapy in EC has been recognized by the NRG and GOG, which sponsored a randomized trial (GOG-0238) in this specific clinical setting.^46^ Given this, we assessed the role of pharmacologic exploitation of intact/wild-type p53 signaling using MDM-2 inhibitors. Both Nutlin-3 and AMG-232 increased p53/p21 signaling and conferred a synergistic effect on parental cells with wild-type *TP53,* particularly at low doses of radiation and MDM-2 inhibitors. Given that the majority of EC (80%) retain wild-type *TP53*, this foundation could lead to genomic-based radio-sensitizing schemes for low-grade EC. Furthermore, we noted that even in the presence of a GOF mutation, MDM-2 inhibitors provide at least an additive effect when given in combination with radiotherapy, presumably leveraging the remaining a wild-type copy. This further broadens the potential application of MDM-2 inhibition in personalized radiation therapy. Further *in-vivo* experimentation is needed to validate these translational findings.

As with most malignancies, alterations in *TP53* confer a poor prognosis. Yet there is emerging data that supports the consideration of not just *TP53* mutation status, but also the variant allelic frequency (VAF), in clinical decision-making. In MDS/AML, patients with a higher VAF *TP53* mutation were shown to have worse overall survival compared patients with lower VAF *TP53* alterations, which had clinical outcomes that approximated those of patients with wild-type *TP53.* Within our the TCGA data, the subgroup of patients with low-VAF *TP53* alterations had outcomes after radiotherapy similar to those of patients with wild-type *TP53*. Notably, the median VAF was not significantly different between the no RT and RT groups (Figure 1E), highlighting that despite the established clinical difference in outcomes the administration of RT is not currently tailored based on *TP53* status. Understanding the molecular and therapeutic impact, as well as the predictive capabilities of VAF *TP53* in EC, may allow treatment personalization.

Taken together, this information supports that an intact p53 signaling pathway may remain functionally relevant in a significant group of EC patients, which could potentially include the 15% of patients with low-VAF mutations. Conversely, this data further supports the notion that most *TP53-*mutated EC patients may benefit from chemotherapy in addition to, or instead, of radiotherapy. We recognize that the TCGA data is retrospective, does not account of different radiation schemes, and has a high degree of variability in recording, thus these results are purely meant to be exploratory and hypothesis generating. Additionally, stratification based on VAF using a 0.5 cutoff is a crude measure of true allelic frequency, as it does not account for gene copy number alteration, ploidy, tumor purity, or clonal enrichment. As this cutoff was demonstrated to be clinically relevant in myeloid leukemias^47^ and would allow us to represent tumors with loss of heterozygosity or homozygous *TP53* mutations, our analysis was conducted using this limit.

In summary, our study provides a biologic rationale behind the observed clinical outcomes of many gynecologic oncology trials; *TP53* alterations confer resistance to radiotherapy in EC. Furthermore, we demonstrate that dominant-negative alleles impact radiation response via abrogation of the canonical p53/p21 pathway and may also interfere with TNF-alpha response. Lastly, we provide both translational data supporting for the investigation of MDM-2 inhibitors as novel radio-sensitizing agents in this disease site, as well as outcomes data that argues for the consideration of its use in both wild-type and low-VAF *TP53*-mutant EC cases. Taken together, our data both supports the established clinical trials regarding p53-aberrant EC, while also advocating for genomics-driven clinical trials that incorporate biologics with radiotherapy in recurrent and early-stage EC.

## Methods

### Cell culture

JHUEM1, JHUEM2, and Hec108, cells were grown in DMEM-F12 media containing 15 mM HEPES and 2.5 mM L-Glutamine (Thermo Fisher, MA) and supplemented with 10% FBS, 100 U/mL Penicillin, 100 μg/mL of Streptomycin. Hec1A, Hec1B, MFE319, and HEK293T cell lines were grown in DMEM media containing 4.5 g/L Glucose, 1 mM Sodium Pyruvate, 4 mM L-Glutamine (Thermo Fisher, MA) supplemented with 10% FBS, 100 U/mL Penicillin, 100 μg/mL of Streptomycin. All cultures were maintained at 37 °C in a humidified 5% CO2 atmosphere. All cell lines were verified by human short-terminal repeat (STR) analysis (Labcorp, NC) and tested to ensure the absence of *Mycoplasma* via qPCR.

### Lentivirus production and infection

Lentivirus containing all mentioned plasmids was generated by transfecting HEK293T cells with the appropriate plasmid along with the psPAX2 (Addgene; Plasmid #12260) and pMD2.G (Addgene; Plasmid #12259) plasmids in a 5:4:1 ratio. Around 18 hours following transfection media was changed, followed by the collection and purification of lentiviral-containing supernatants over the next two days. Centrifugation and passage through a 40 uM filter were used for purification. Cell lines were infected with these viruses at an MOI > 0.8 in media supplemented with 5 ug/ml Polybrene.

### Exon-intron junctional CRISPR

sgRNAs that target the exon 5-intron (5.1) and exon 6-intron (6.1) junctions of *TP53* were designed with the Synthego CRISPR web design tool (https://www.synthego.com/products/bioinformatics/crispr-design-tool/). Non-targeting control (NTC) sgRNAs were also designed. For each sgRNA, two complementary oligos with BsmBI digestion overhang sequences were designed. Sequences of sgRNAs can be found in **Supplementary Table 3.** plentiCRISPRv2-Blast plasmid (Addgene: Plasmid #83480), which contains the Cas9 coding sequence and a cloning site for sgRNA, was digested with BsmBI, followed by annealing of sgRNAs and ligation into the pLentiCRISPRv2-Blast plasmid as in the Zhang Lab protocol given at http://www.genome-engineering.org/gecko, as well as on the website for the original pLentiCRISPRv2 plasmid (Addgene #52961).^48,49^ Plasmids were transformed into NEB Stable bacteria (New England Biolabs, MA) and colonies were screened for sgRNA inserts by Sanger sequencing of the ligation site. After virus infection, cells were selected and maintained in the presence of 5 μg/mL blasticidin (Thermo Fisher, MA). Monoclonal cells were isolated from a polyclonal pool of transduced stable cells using the limiting dilution assay. Protein lysates were obtained from each clone and *TP53* KO was confirmed by Western Blot.

### Variant-expressing plasmid generation

Variant-expressing lentiviral plasmids were generated by utilizing PCR-based site-directed mutagenesis, followed by bacterial recombination and transformation. First, 5’ and 3’ fragments containing incorporated mutated gene ORF sequences were amplified by PCR from the pDONR223-*TP53* WT plasmid (Addgene; Plasmid #82754). In this case, the 5’ and 3’ flanking PCR primers were located at the attL1 and attL2 sites, with the internal primers containing incorporated mutated sequences. Sequences of PCR primers can be found in **Supplementary Table 4.** Fragments were then transferred to the destination vector pLEX306 (Addgene; Plasmid #49391) by LR reaction with LR Clonase II enzyme (Invitrogen, CA), followed by transformation into competent cells. The overlapping fragment overhangs at the mutation site were repaired by endogenous bacterial repair. Inserted sequences and incorporated mutation were verified by restriction digestion with BsrGI and Sanger sequencing of ORF, respectively.

### Complementation

Wild type and variant containing ORFs of *TP53* were cloned into pLEX306 and integrated into the respective CRISPR KO or non-target control (NTC) cell lines. After virus infection, cells were selected and maintained in the presence of 1 μg/mL puromycin.

### Antibodies and reagents

Anti-p53 (#9282, 1:3000), anti-p21 Waf1/Cip1 (clone 12D1, #2947, 1:3000), anti-Vinculin (clone E1E9V, #13901, 1:5000) were from Cell Signaling Technology. Goat anti-rabbit antibody linked to horseradish peroxidase (HRP) was used as the secondary antibody (Santa Cruz Biotechnology, CA).

### Western blot analysis

Cells were pretreated with 10 uM Nutlin-3 in 0.1% DMSO for 18 hours to enhance the protein signal to a detectable level for comparison. Cells for dose-response analyses were treated with an appropriate dose of 0.05% DMSO. Whole-cell lysates were prepared using M-PER lysis buffer (Thermo Fisher, MA) and clarified by centrifugation. Proteins were separated by SDS–PAGE on 4-12% Bis-Tris gradient gels (Thermo Fisher, MA) and transferred onto 0.2 μM nitrocellulose membranes (Bio-Rad, CA). Membranes were blocked and incubated in primary antibody for 2 hours at room temperature, followed by washing and incubation for 1h with secondary antibody. Blots were developed with the ECL Plus chemiluminescence system (Amersham/GE Healthcare, UK).

### High-content radiation assay

Cells were plated using a Multidrop Combi liquid handler (Thermo Fisher, MA) at multiple cell densities in white 96-well plates (Corning, NY) for each cell line. Cells were plated in 6 replicate wells for each cell density in triplicate. Plates were irradiated and at 9 days post-irradiation, media were aspirated and 50 μL CellTiter-Glo^®^ reagent (50% solution in PBS) (Promega, WI) was added to each well. Relative luminescence units (RLU), proportional to the amount of ATP present, were measured using a Synergy H1 (Biotek, IL) luminescence plate reader. The luminescence signal was plotted as a function of cell density and a cell density within the linear range for luminescence was selected to generate integral survival for each cell line.^35^ Cells were treated with γ-radiation delivered at 0.92 Gy/min with a ^137^Cs source using the Mark 1 Irradiator (Shepherd and Associates, CA). The area under the curve (AUC) was estimated using a trapezoidal approximation in GraphPad Prism software (Version 8.4.3, GraphPad, CA). Radiation doses of 0, 2, 4, 6, and 8 Gy were given and the fraction surviving was determined by normalizing each RLU value to the 0 Gy value. These data were then used to generate an AUC value for each cell line.

### Drug dose-response assays

For drug dose-response experiments, cells were plated as given above and treated with either 0.5% DMSO (Control), or 7 different doses of the drug in 0.5% DMSO. For Nutlin-3, doses of 0.1, 0.5, 1, 5, 10, 50, 100 uM were used and 0.01, 0.05, 0.1, 0.5, 1, 5, 10 nM for AMG-232. Cells were treated for 24 hours, followed by replacement with drug-free media and incubation for 9 days. Cells were then lysed with CellTiter-Glo reagent and RLU values were obtained as given above. Fraction surviving was determined by normalizing each RLU value to either the 0 Gy value, or the maximum value, whichever was greater. Four-parameter sigmoid curves were fitted to the dose-response data and IC50 values were determined using GraphPad Prism software (Version 8.4.3, GraphPad, CA).

### Drug radio-sensitization assays

For drug radio-sensitization experiments, cells were plated and treated with drug doses as given above. Following replacement with drug-free media, radiation doses of 0, 0.5, 1, 2, 4, and 8 Gy were given, followed by incubation for 9 days. For JHUEM2 and Hec108, cells in plates receiving 2, 4, and 8 Gy of radiation were plated at twice the density to allow accurate RLU signal detection and comparisons. Fraction surviving was determined by normalizing each RLU value to the Untreated (0 uM, 0 Gy) value. Fraction surviving values at 2, 4, and 8 Gy were halved to correct for higher plating density. The mean values for three replicates were taken and all were divided by the maximum signal. These data were used to perform synergy analysis. Briefly, the Highest Single Agent (HSA) additivity model was used to generate predicted additive response curves for the JHUEM2 and Hec108 data. The Loewe additivity model was used for Hec1B, as the data did not meet the requirements for HSA. The actual data was then compared and synergy scores generated based on the deviation of each point from the model.

### RNA Isolation and RNA Seq

For RNA isolation from cell lines, cells were first lifted off plates, washed with 1X PBS, and pelleted by centrifugation. These pellets were then lysed and RNA extracted using the AllPrep DNA/RNA Mini Kit (Qiagen, MD). RNA Seq analysis was performed by the in-house Genomics Core Facility at The Cleveland Clinic. Briefly, the total RNA samples were enriched for mRNAs, followed by library preparation and fragmentation using the Illumina TruSeq Kit (Illumina, CA). Paired-end 100 bp sequencing reads were generated using the Illumina NovaSeq 6000 machine (Illumina, CA). RNA-Seq read pre-processing was done using *fastp* with low complexity filters set and trimmed from both 3’ and 5’ ends.^50^ Reads were then aligned to human genome gencode v40 using STAR aligner in gene quantification mode with 2 pass enabled.^51^ Differential expression was performed using R package *edgeR.*^52^ Pathway enrichment was completed using Hallmark Gene Set from MsigDB and p-values were calculated via hypergeometric distribution.^53^

### Survival Analysis

Clinical data was obtained from cbioportal.org and curated from the uterine cancer TCGA Firehose Legacy, Nature 2013, and PanCancer Atlas datasets.^54,55^ All patient identifiers were matched to collate clinical and genomic markers for analysis. Duplicate samples were removed. Only patients with tumor genomic data (*TP53* status, allelic frequency, tumor mutational burden, fraction genome altered, biologic cluster) and clinical data (age at diagnosis, race, FIGO stage, tumor histologic grade, adjuvant radiotherapy, progression-free survival) were included in the final dataset for analysis. Patients with multiple *TP53* mutations or stage IV disease were excluded, as well as those receiving adjuvant chemotherapy. Consistent with the TCGA analysis, a percent genome altered greater than 0.2 was considered copy-number high. ^37^ Similarly, VAF was stratified using a 0.5 allelic frequency cutoff Kaplan-Meier and Cox proportional hazards analyses were performed to assess for impact on progression-free survival (PFS). Multivariable analyses were performed on clinical/genomic variables with significance on univariate analysis.

### Graphs and statistical analyses

Statistical analyses for TCGA clinical data were performed using R 4.2.1 and R Studio 2021.09.1.^56^ For all analyses in R, data in the form of Excel tables were read into R using the readxl package.^57^ For Kaplan-Meier analysis, Cox proportional hazards statistical analysis was performed and the results are those quoted. Survival analyses were performed using R package survival^58,59^and survival graphs were generated with survminer.^60^ Graphs were generated and statistical analyses were performed for drug radiosensitization studies using the BIGL package.^61^ All graphs for clinical data were generated in R using the ggplot2 and RColorBrewer packages.^62,63^ Statistical analyses for cell line data were performed using the GraphPad Prism software (Version 8.4.3, GraphPad, CA). Graphs for cell line data were generally made using GraphPad Prism software, except those for Figures 2A, 2B, and 3B, which were generated using the R packages for clinical data above, as well as the ggbeeswarm package.^64^ All clinical data summary tables were generated using the gtsummary package.^65^ R code for graphs, tables, and statistical analyses are available on GitHub at the following address (www.github.com/petty1284/TP53-and-Endometrial-Cancer).

## Acknowledgements

We wish the acknowledge The Laura J. Fogarty Endowed Chair in Women’s Health Institute for Uterine Cancer and thank them for their unwavering support of our mission.

