## Supplementary Figures and Tables for "Leveraging the p53 signaling pathway as a radio-sensitization strategy in endometrial cancer"

### Supplementary Materials

### Supplementary Figure 1.

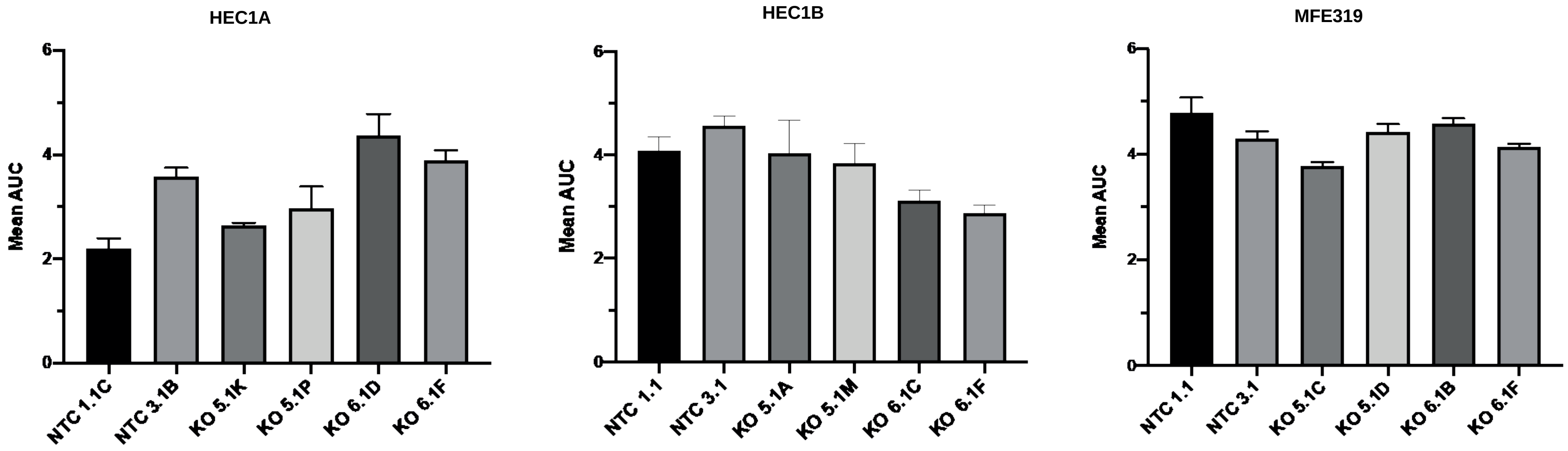


TP53 knockout via CRISPR/Cas9 in three EC cell lines with GOF/DN alleles and high VAF (presumed loss-of-heterozygosity) does not impact radiation response.

### Supplementary Figure 2

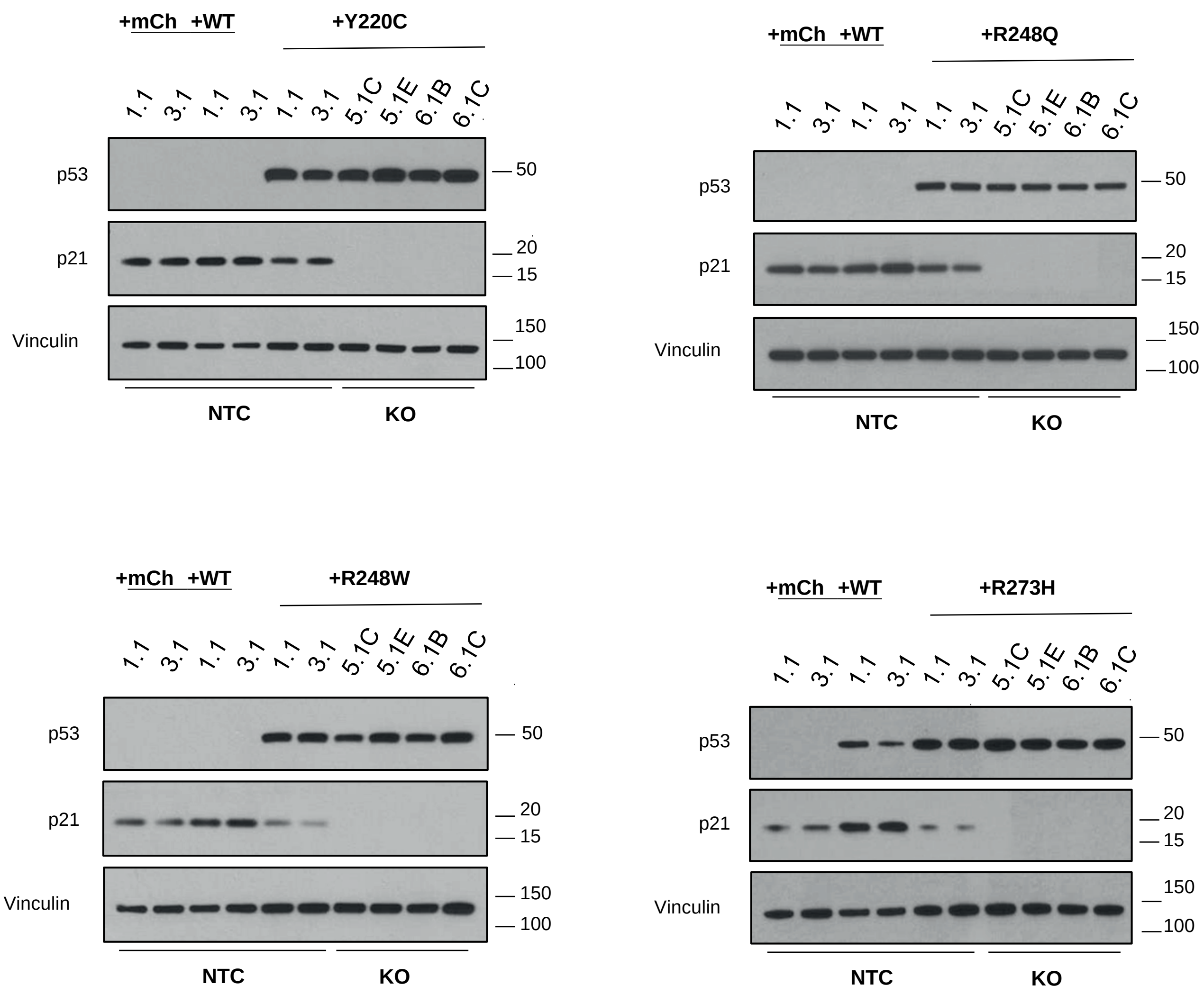


Western-blot analysis of 4 alternative TP53 alleles reveal abrogation of p21 signaling by the mutant allele in the presence of two wild-type alleles. As seen in the knock-out monoclonal lines (5.1C/E and 6.1B/C) expression of the mutant allele alone does not impact p21 signaling but confers significant accumulation of p53.

### Supplementary Figure 3.

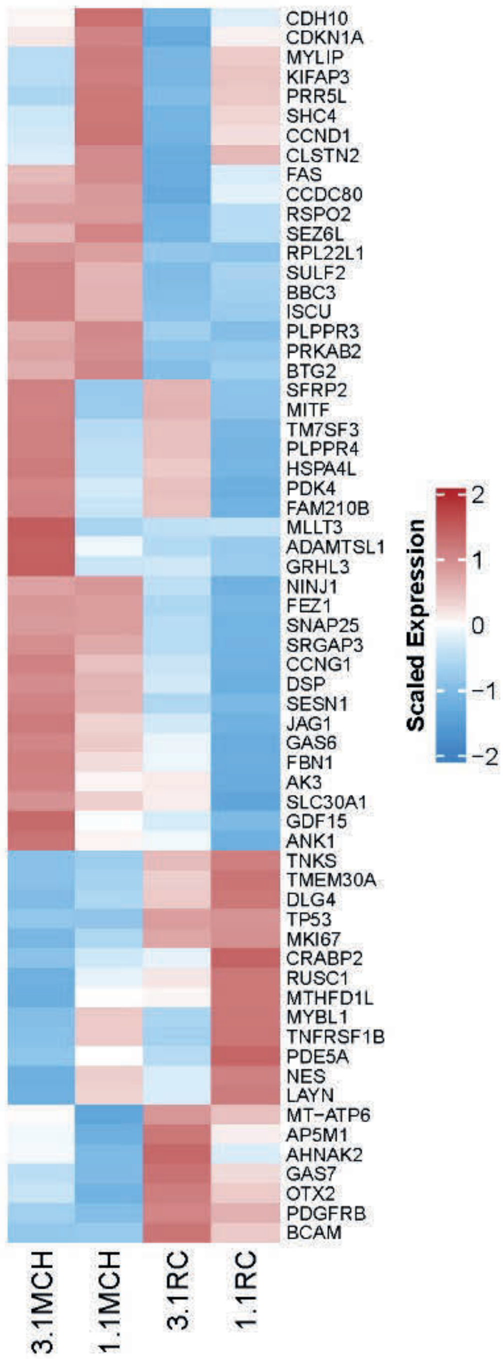


Heat-map of differential gene expression. In this experiment, two NTC (1.1 and 3.1) monoclonal isolates were transfected with mcherry and an R273C allele. Radiation was administered (2Gy) and RNA extracted 24 hours later. Differential gene expression between the lines with only wild-type alleles and those with wild-type + R273C can be appreciated. Notably TP53 expression is confirmed in our 1.1RC and 3.1RC constructs, confirming increased expression via our PGK promoter.

### Supplementary Figure 4.

A.

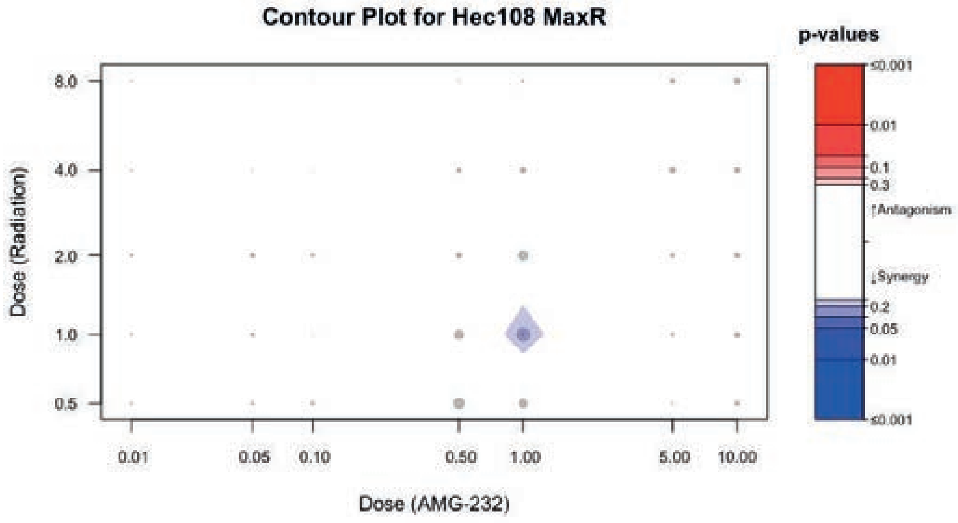


B.

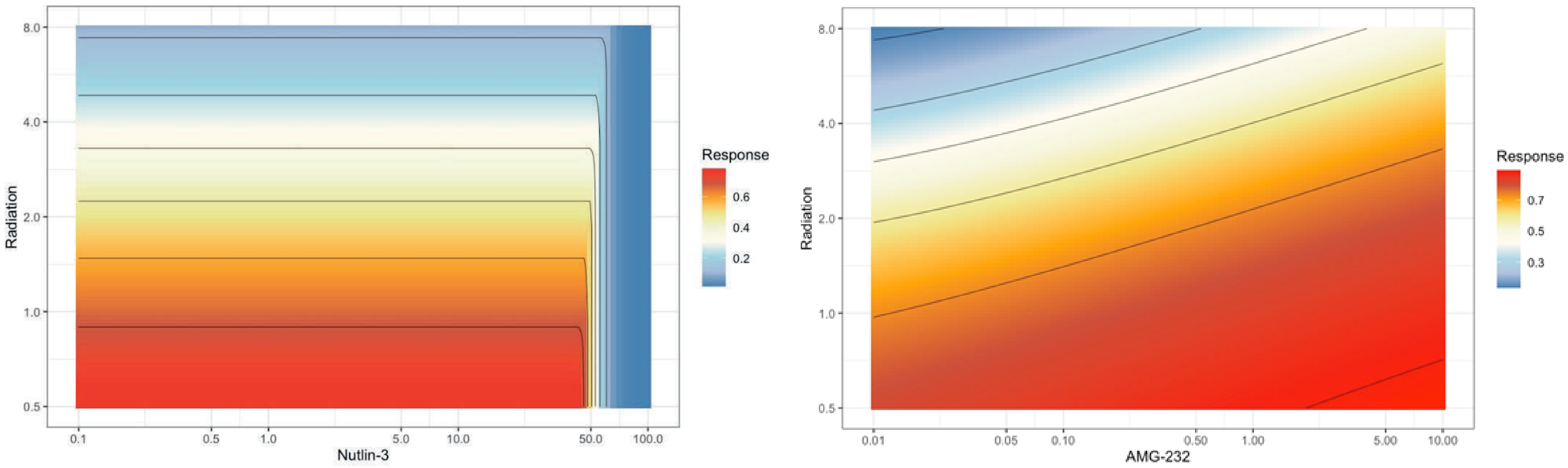


A) Contour plot of HEC108 and AMG-232 demonstrating weak synergy at 1nM and 1Gy dosing. **B)** Isobologram of Hec1B using radiation (y-axis) and drug concentration (x-axis). As noted there is no effect with the addition of Nutlin-3 to radiotherapy and a suggestion of an antagonistic effect at higher doses with AMG-232.

### Supplementary Figure 5.

A.

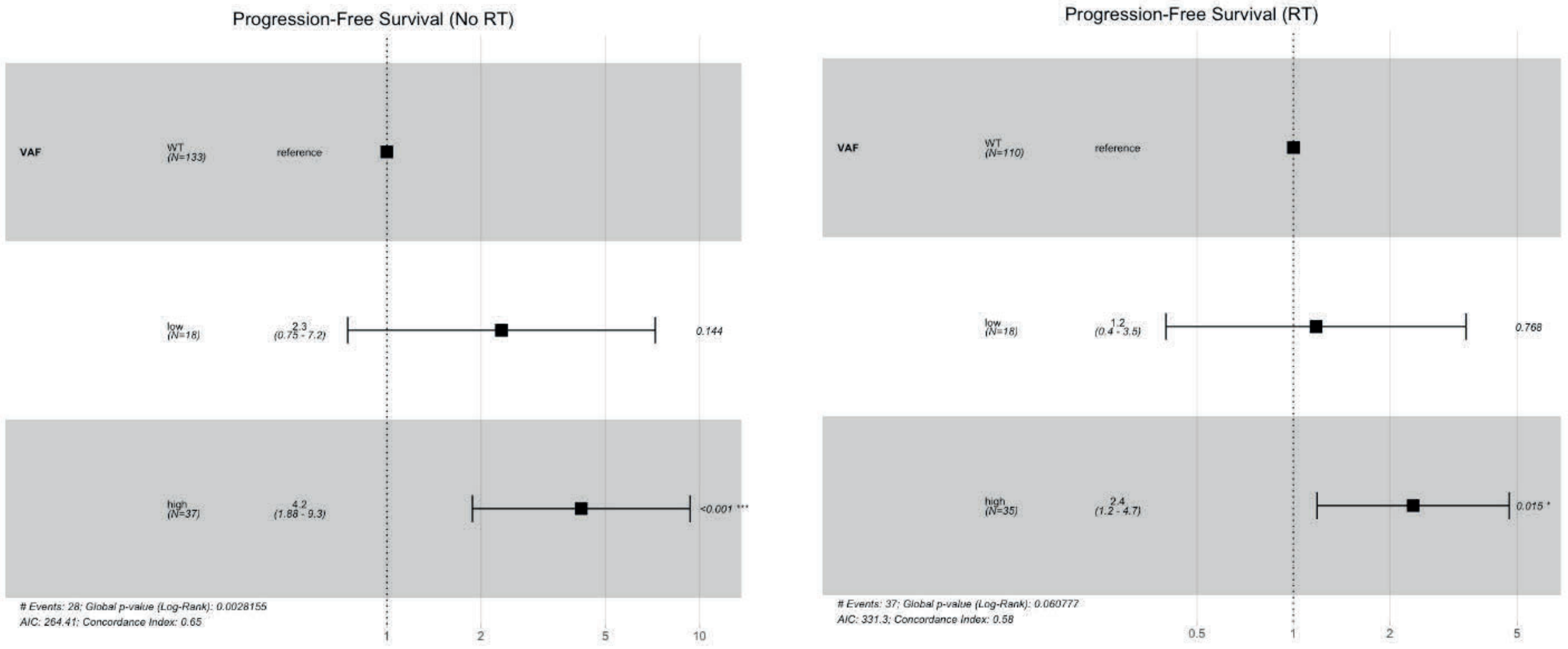


B.

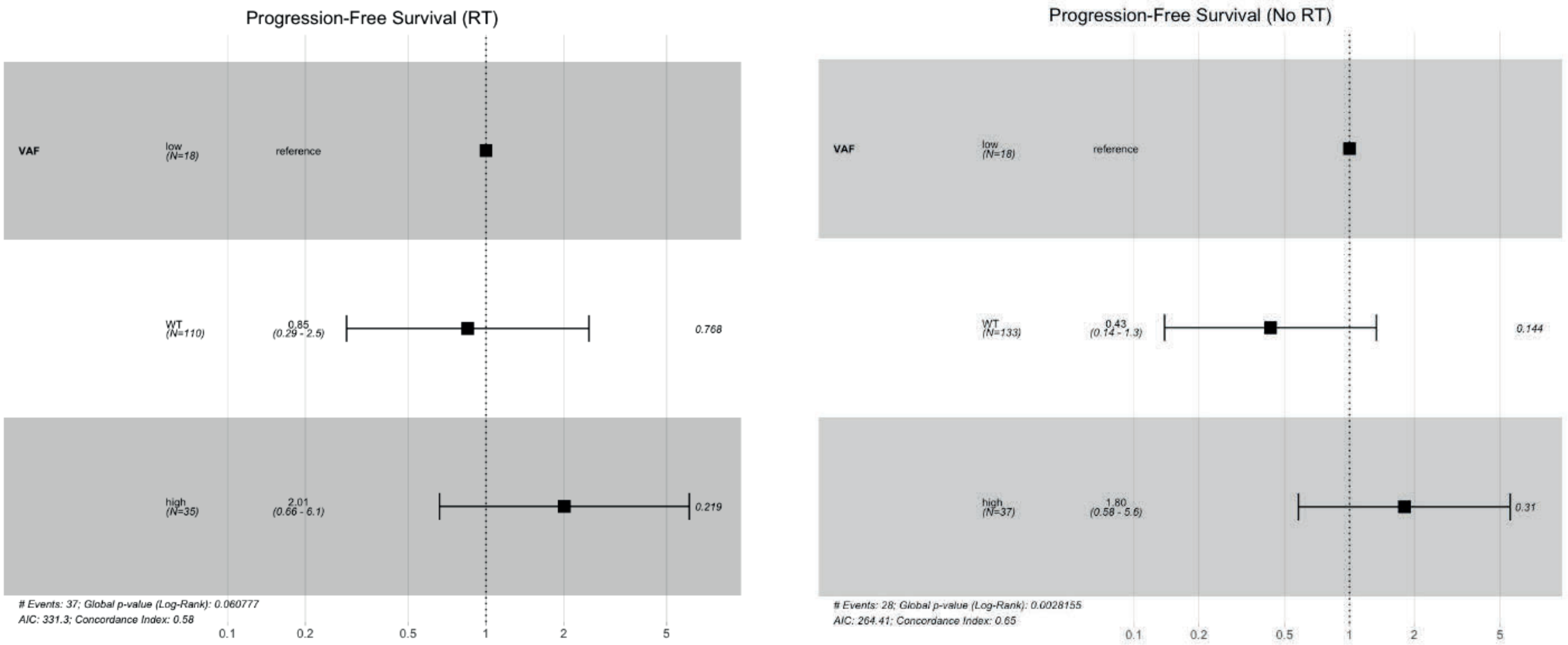


COX Regression analysis using wild-type **(A)** and low-VAF (**B**) as the reference for analysis. As noted, risk of progression is higher in patients with high-VAF tumors compared to wild-type patients. While no difference is noted when comparing low-VAF to wild-type patients without radiation, the hazard ratio shift with the receipt of radiotherapy suggests a notable response. Using low-VAF as the reference, no statistically significant differences are observed.

### Supplementary Table 1.

A.

| Characteristic | Radiotherapy Received: No, N = 188[1] | Radiotherapy Received: Yes, N = 163[1] | p-value[2] |
|---|---|---|---|
| **Age at Diagnosis** | 64.00 (11.30) | 63.00 (10.93) | 0.3 |
| **Total Mutation Count** | 63.00 (1,942.69) | 112.00 (2,393.62) | 0.006 |
| **Fraction Genome Altered** | 0.05 (0.22) | 0.05 (0.21) | 0.8 |
| **TCGA Cluster** | | | 0.4 |
| Not Classified | 2 / 188 (1.1%) | 4 / 163 (2.5%) | |
| UCEC_CN_HIGH | 55 / 188 (29%) | 42 / 163 (26%) | |
| UCEC_CN_LOW | 68 / 188 (36%) | 49 / 163 (30%) | |
| UCEC_MSI | 50 / 188 (27%) | 56 / 163 (34%) | |
| UCEC_POLE | 13 / 188 (6.9%) | 12 / 163 (7.4%) | |
| **FIGO Stage** | | | 0.5 |
| I | 134 / 188 (71%) | 108 / 163 (66%) | |
| II | 15 / 188 (8.0%) | 18 / 163 (11%) | |
| III | 39 / 188 (21%) | 37 / 163 (23%) | |
| **Histologic Grade** | | | 0.003 |
| 1 | 53 / 188 (28%) | 22 / 163 (13%) | |
| 2 | 50 / 188 (27%) | 46 / 163 (28%) | |
| 3 | 85 / 188 (45%) | 95 / 163 (58%) | |
| **P53 Mutation** | 55 / 188 (29%) | 53 / 163 (33%) | 0.5 |

[1] Median (SD); n / N (%)

[2] Wilcoxon rank sum test; Fisher's exact test; Pearson's Chi-squared test

B.

| Characteristic | Progression-Free Survival: HR[1] | 95% CI[1] | p-value |
|---|---|---|---|
| **sCNA** | | | |
| *Low* | — | — | |
| *High* | 3.04 | 1.86, 4.95 | <0.001 |
| **Age** | | | |
| *< 50* | — | — | |
| *50-70* | 2.71 | 0.66, 11.1 | 0.2 |
| *> 70* | 2.12 | 0.48, 9.26 | 0.3 |
| **p53** | | | |
| *WT* | — | — | |
| *VAF Low* | 1.64 | 0.75, 3.56 | 0.2 |
| *VAF High* | 3.26 | 1.93, 5.50 | <0.001 |
| **Radiotherapy** | | | |
| *No* | — | — | |
| *Yes* | 1.59 | 0.97, 2.60 | 0.065 |
| **Stage** | | | |
| *I* | — | — | |
| *II* | 1.07 | 0.42, 2.73 | 0.9 |
| *III* | 2.64 | 1.58, 4.40 | <0.001 |
| **Grade** | | | |
| *1* | — | — | |
| *2* | 2.17 | 0.89, 5.27 | 0.088 |
| *3* | 2.80 | 1.26, 6.25 | 0.012 |

[1] HR = Hazard Ratio, CI = Confidence Interval

C.

| Characteristic | Progression-Free Survival: HR[1] | 95% CI[1] | p-value |
|---|---|---|---|
| **sCNA** | | | |
| *Low* | — | — | |
| *High* | 2.03 | 0.96, 4.28 | 0.063 |
| **Stage** | | | |
| *I* | — | — | |
| *II* | 0.79 | 0.31, 2.05 | 0.6 |
| *III* | 2.10 | 1.24, 3.56 | 0.006 |
| **Grade** | | | |
| *1* | — | — | |
| *2* | 1.84 | 0.75, 4.52 | 0.2 |
| *3* | 1.34 | 0.53, 3.37 | 0.5 |
| **p53** | | | |
| *WT* | — | — | |
| *VAF Low* | 1.14 | 0.48, 2.71 | 0.8 |
| *VAF High* | 1.65 | 0.74, 3.70 | 0.2 |

[1] HR = Hazard Ratio, CI = Confidence Interval

(A) Demographics table stratified by receipt of radiotherapy. Univariate (B) and multivariable (C) analysis of progression-free survival.

### Supplementary Table 2.

A.

| Term | Overlap | P-value | Adjusted P-value | Odds Ratio | Combined Score | Genes |
|---|---|---|---|---|---|---|
| **p53 Pathway** | **9/200** | **3.59E-06** | **1.26E-04** | **8.283096** | **103.8415782** | **CDKN1A;BTG2;SESN1;NINJ1;CCNG1;FAS;TM7SF3;TP53;BLCAP** |
| **Apoptosis** | **6/161** | **4.44E-04** | **0.007778** | **6.639215** | **51.24539677** | **CDKN1A;BTG2;CCND1;GADD45B;PLPPR4;FAS** |
| **TNF-alpha Signaling via NF-kB** | **6/200** | **0.001375** | **0.016047** | **5.294039** | **34.88214092** | **CDKN1A;BTG2;CCND1;GADD45B;NINJ1;IER2** |
| Mitotic Spindle | 3/199 | 0.119967 | 0.422972 | 2.553139 | 5.414022257 | RICTOR;KIFAP3;KIF15 |
| Hypoxia | 3/200 | 0.121286 | 0.422972 | 2.54005 | 5.358499671 | CDKN1A;HOXB9;CCNG2 |
| G2-M Checkpoint | 3/200 | 0.121286 | 0.422972 | 2.54005 | 5.358499671 | CCND1;MKI67;KIF15 |
| Estrogen Response Early | 3/200 | 0.121286 | 0.422972 | 2.54005 | 5.358499671 | SOX3;CCND1;MYBL1 |
| Myogenesis | 3/200 | 0.121286 | 0.422972 | 2.54005 | 5.358499671 | CDKN1A;GADD45B;SCHIP1 |
| mTORC1 Signaling | 3/200 | 0.121286 | 0.422972 | 2.54005 | 5.358499671 | CDKN1A;BTG2;CCNG1 |
| E2F Targets | 3/200 | 0.121286 | 0.422972 | 2.54005 | 5.358499671 | CDKN1A;MKI67;TP53 |

A) Hallmark pathway analyses from differential gene expression obtained 24 hours after radiotherapy exposure. See supplementary data for differential gene expression.

### Supplementary Table 3.

| Target | Target Sequence | F oligo (BsmBI site in red) | R oligo (BsmBI site in red) |
|---|---|---|---|
| NTC 1.1 | GTATTACTGATATTGGTGGG | CACCGGTATTACTGATATTGGTGGG | AAACCCCACCAATATCAGTAATACC |
| NTC 3.1 | TCAACCCCAGCGCACCGTTG | CACCGTCAACCCCAGCGCACCGTTG | AAACCAACGGTGCGCTGGGGTTGAC |
| TP53 5.1 | TGAGGGCAGGGGAGTACTGT | CACCGTGAGGGCAGGGGAGTACTGT | AAACACAGTACTCCCCTGCCCTCAC |
| TP53 6.1 | GTTGCAAACCAGACCTCAGG | CACCGGTTGCAAACCAGACCTCAGG | AAACCCTGAGGTCTGGTTTGCAACC |

CRISPR/Cas9 guide-RNA sequences. A designed 5.1 targets exon 5 and 6.1 targets exon 6.

### Supplementary Table 4.

| Name/mutation | F primer (mutated site underlined) | R primer (mutated site underlined) |
|---|---|---|
| attL flanking sites | (L1) CGTTGTAAAACGACGGCCAGTC | (L2) GCCAGGAAACAGCTATGACCATG |
| TP53 R273H | AGCTTTGAGGTGCATGTTTGTGCCTGT | ACAGGCACAAACATGCACCTCAAAGCT |
| TP53 R273C | AGCTTTGAGGTGTGTGTTTGTGCCTGT | ACAGGCACAAACACACACCTCAAAGCT |
| TP53 R248Q | GGCGGCATGAACCAGAGGCCCATCCTC | GAGGATGGGCCTCTGGTTCATGCCGCC |
| TP53 R248W | GGCGGCATGAACTGGAGGCCCATCCTC | GAGGATGGGCCTCCAGTTCATGCCGCC |
| TP53 Y220C | GTGGTGGTGCCCTGTGAGCCGCCTGAG | CTCAGGCGGCTCACAGGGCACCACCAC |

*TP53* variant site-directed mutagenesis primer sequences with nucleotide substitutions.
